# Fine-scale oceanographic processes shape marine biodiversity patterns in the Galápagos Islands

**DOI:** 10.1101/2024.03.06.583537

**Authors:** Luke E Holman, Diana A. Pazmiño, Shyam Gopalakrishnan, Alexander Forryan, Alex R. Hearn, Alberto C. Naveira-Garabato, Marc Rius

## Abstract

Uncovering the drivers that shape biodiversity patterns is critical to understand ecological and evolutionary dynamics. Despite evidence that biodiversity composition is influenced by processes at different spatial scales, little is known about the role of fine-scale oceanographic processes on the structure of marine communities. This is particularly important in biodiversity hotspot regions, where small changes in environmental conditions may lead to substantial changes in species composition. We combined ocean modelling and 12S environmental DNA (eDNA) metabarcoding, targeting teleost and elasmobranch species, to explore if oceanographic processes influenced biogeographic patterns around the biodiverse Galápagos Islands. We first detected significant differences in eDNA-measured community structure across the archipelago’s diverse seascape. We found no significant relationship between Lagrangian particle tracking metrics and nektonic biodiversity, and thus developed a novel metric to measure the cumulative seawater flow resistance between pairs of geographic sites. This metric explained a significant proportion of variation in eDNA-measured beta dissimilarity between sites, comparable in influence to important abiotic drivers, such as temperature and geographic distance between sites. Cumulatively, our results indicate that marine communities are particularly sensitive to changes in local current systems, and suggest that fine-scale oceanographic processes may have an underappreciated role in structuring marine communities globally.

## Introduction

Spatial patterns of biodiversity are profoundly influenced by physical factors, such as oceanic currents and geographic barriers, with direct consequences on species distributions and community structure ^1–3^. Indeed, research has shown that planktonic communities increase in similarity proportionally to oceanographic connectivity ^4–6^, with distance travelled along currents having the same effect on beta diversity as geographic distance on land (so called distance-decay relationships) ^7,8^. Despite decades of research to understand how oceanography shapes population connectivity, there are very few studies that explore the importance of local or submesoscale (horizontal scales ≪ 100 km) changes in ocean currents relative to other variables (such as temperature) in controlling marine biodiversity patterns ^9,10^. Furthermore, we currently lack insight into whether submesoscale ocean currents structure plankton biodiversity across ecosystems ^11^, and to what extent such currents affect free-swimming non-planktonic (nektonic) communities. This is particularly important in biodiversity hotspot regions, where small changes in environmental conditions may lead to substantial biodiversity changes ^12,13^.

Ocean currents are central in shaping the distribution and movement of species that complete part or all of their life cycle adrift in the ocean. Fine-scale ocean currents may shape the distribution of nektonic species because: (i) a substantial proportion of nektonic organisms have planktonic early-life history stages ^14^; (ii) plankton and nekton are tightly connected through food webs ^15,16^; and (iii) nektonic organisms tend to track thermal optima in current systems ^17^. Conversely, the distribution of some nektonic species may not be related to currents, as these species can migrate thousands of kilometres moving across current systems ^18^ to exploit resources that can be affected by ocean circulation ^19,20^. Some studies have examined how currents affect whole community diversity ^5,9,10^, and seascape genetics research has examined the effect of current systems on population genetics across different ocean systems ^21,22^. All these studies use some form of Lagrangian particle modelling as a surrogate for the current systems encountered by organisms. This modelling approach tracks ocean currents by virtually releasing particles into an oceanographic model, estimating the likelihood of a particular particle reaching a location or measuring the time taken for a given journey. In sufficiently mixed systems, these metrics work well, but when sites are only very infrequently connected (for example, a complex island system with a prevailing current direction), Lagrangian particles are not able to provide accurate measures of the differences between currents. Moreover, some organisms cross vast ocean distances, migrating through current systems to reach resources ^19,20^. In these cases Lagrangian particle tracking may not provide any estimate of the current forces encountered across these migrations, as particles simply are unlikely to pass through several current systems. To comprehensively understand the effect of currents on marine communities, Langragian particle tracking may require additional complementary metrics to capture how current systems affect biodiversity patterns.

In recent years, the use of high-throughput sequencing to analyse DNA isolated from the environment (often called environmental DNA or eDNA) has become common practice, and is now an established approach for producing whole-community data ^23,24^. Our understanding of marine biodiversity is being revolutionised through eDNA surveys, with research uncovering previously undocumented global patterns ^2,5^, revealing previously undescribed taxonomy ^25^ and, most recently, reconstructing long-dead marine taxa and biodiversity from ancient eDNA ^26,27^. Despite all these advances, marine eDNA studies rarely integrate ocean circulation into their analyses ^5,28,29^, and non-eDNA studies in community ecology have only explored the link between eDNA patterns and ocean currents with a relatively small subset of taxa^9,10^. There is therefore a pressing need to understand the potential role of ocean flows on biodiversity patterns, considering a wide array of both planktonic and non-planktonic organisms.

Ocean currents play a central role in shaping the diversity of marine life of the Galápagos archipelago ^30^. Located in the Eastern Pacific, the Galápagos Islands are an iconic biodiversity hotspot, rising at the confluence of the South Equatorial, Cromwell and Humboldt current systems ^31^. The remarkable biodiversity of the region depends on the wind-driven upwelling of cool, nutrient rich waters ^32^. This upwelling system results in abundant phytoplankton on the western side of the archipelago ^33^, particularly the islands Isabela and Fernandina (see Fig. S1), forming the basis of a uniquely productive region characterised by local bioregionalisation ^30^, and elevated marine endemism^34^, biodiversity and functional diversity ^35^. Galápagos fish communities are remarkably diverse, displaying the highest functional diversity at any given species diversity globally ^36^, warranting further study into the patterns and drivers that maintain such communities. Despite substantial research efforts into understanding the marine biodiversity of the Galápagos ^30,34,35^, we still only have a handful of studies that use eDNA to understand the biodiversity of the region ^37–39^, none of which aim to explicitly model the effect of currents on marine communities.

Here, we elucidate the effect that ocean currents have on marine community structure across the waters surrounding the Galápagos islands. We first use eDNA metabarcoding of seawater samples collected from across the archipelago to detect spatial patterns of teleost fish and elasmobranch biodiversity. Subsequently, we model the ocean circulation at high (submesoscale-permitting) resolution and infer the effect of eDNA decay ^40,41^, using Lagrangian particle tracking to better understand the detected patterns of nektonic biodiversity. We then develop a metric that describes local current systems from ocean model-generated data, motivated by the omission of ocean flow pathways in Lagrangian-based metrics. Finally, we assess the comparative effects of ocean currents and sea surface temperature on community dissimilarity.

## Results

### Galápagos fish biodiversity

Metabarcoding of eDNA water samples collected from sites across the Galápagos (Fig. 1a, Table S1) produced a fish (teleost and elasmobranch) dataset containing 551 amplicon sequence variants (ASVs) of which 66 could be assigned to species level, 216 to Genus, 167 to Family, and 99 above Family level. Read numbers and diversity in negative control samples were typical for eDNA metabarcoding datasets ^24^ (full details provided in Supplementary Text 1).

**Fig. 1.**
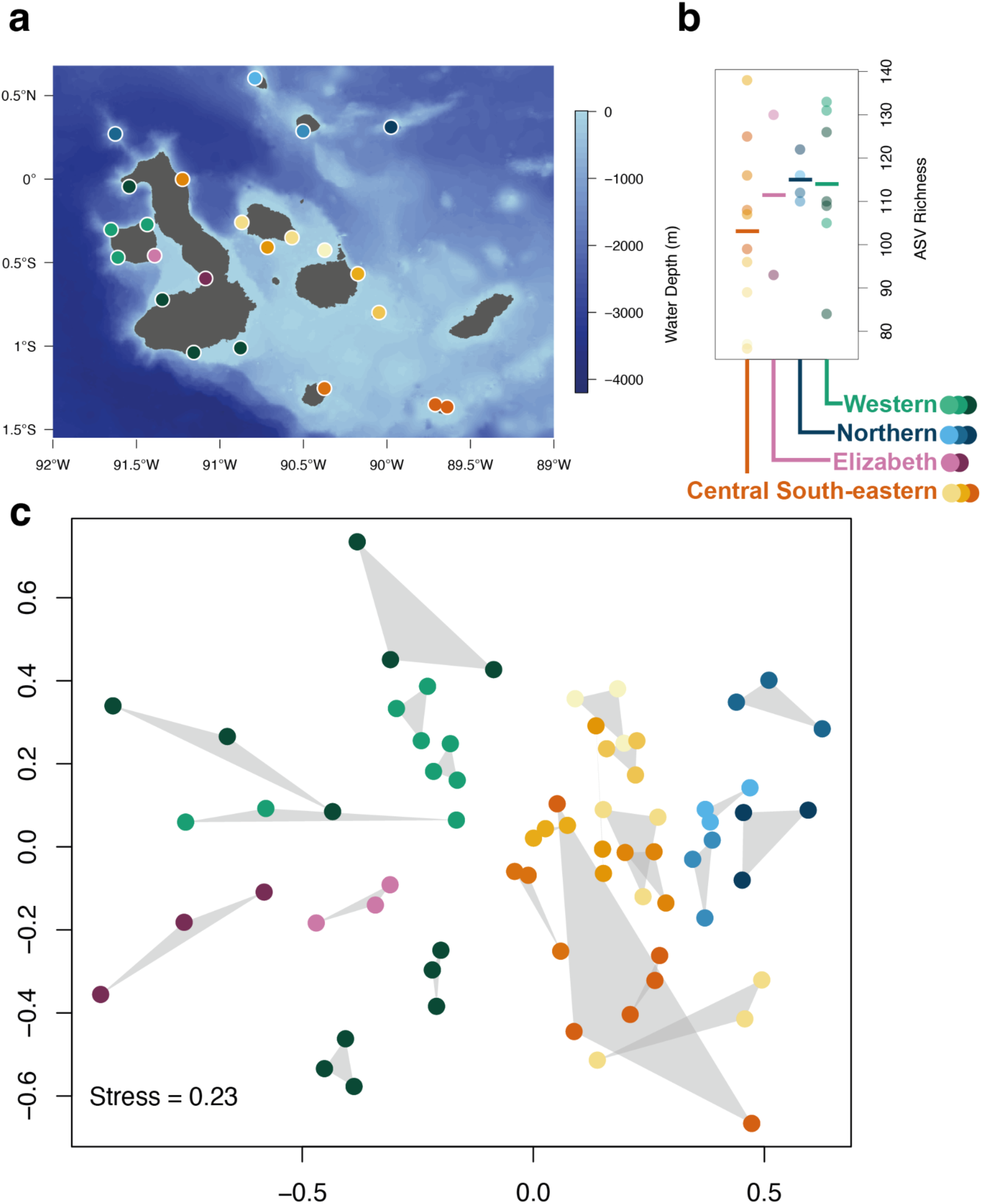
a) Map of the Galápagos islands indicating the sampling sites (marked as dots) and depth (blue colour gradient). b) ASV richness across the sampling sites grouped by four of the bioregions and averaged over field replicates, with the mean value indicated by a solid horizontal line. c) Non-metric multidimensional scaling based on Jaccard dissimilarity of community composition among sampling sites. Each point represents a single field sampling replicate, with the three replicates per site joined by a grey convex hull. In all plots, point colour indicates bioregions from ^30^ as indicated in b), with individual sites coloured with different shades.

Fish communities clustered in the nMDS ordination (Fig. 1c) in agreement with previously reported bioregions ^30^. Specifically, the Elizabeth bioregion clustered within the Western bioregion, and both bioregions were separated from the Northern and Central South-eastern bioregions. In contrast with previous findings Roca Redonda (the most north-westerly site in Fig. 1a and Fig. S1) clustered with sites from the Northern bioregion (top right islands in Fig. 1c), and separated from sites from the Western bioregion.

There was a statistically significant overall difference among bioregions (PERMANOVA F_3,19_ = 2.08, p < 0.001), with pairwise tests showing significant results (p < 0.01) among all bioregions except for the Elizabeth bioregion, which was not significantly (p > 0.05) different to any other bioregion (full model outputs in Table S2). Pairwise tests for significant differences in multivariate dispersion between bioregions (PERMDISP procedure) indicated that only the Elizabeth bioregion had significantly different multivariate dispersion compared to the Central South-eastern and Western bioregions (p < 0.01 in both cases, see Table S1 for full model output) suggesting less within-bioregion variation for the relatively smaller area covered by this bioregion. A one-way ANOVA indicated no significant differences (F_3,19_ = 0.72, p > 0.05) in mean ASV richness among bioregions (Fig. 1b).

### Describing Galápagos currents

Lagrangian particle tracking was used to estimate the current-derived connectivity between pairs of sites. A linear regression indicated that the log_10_ transformed minimum drift time between sites and the log_10_ transformed fraction of particles reaching a given site from the release site were strongly inversely correlated (F_1,241_ = 499.6, R^2^ = 0.674, p < 0.001, see Fig. S2). As these metrics contain overlapping information, we used the minimum drift time in subsequent analyses. The mean drift time across all site-site journeys was 16.8 ± 11.7 (s.d.) days, with a maximum of 54.6 days and a minimum of 0.6 days. Minimum drift time could be computed for 48% of possible journeys, as 52% of journeys could not be completed due to prevailing current direction. Given that fish move against the prevailing current ^18–20^, and to derive a metric that produced a value for paths between all site pairs, we developed a novel metric to describe the current encountered along a seascape journey. We named this new metric oceanographic resistance (Fig. 2), as it describes the average current direction and magnitude between two marine sites for a given journey direction.

**Fig. 2.**
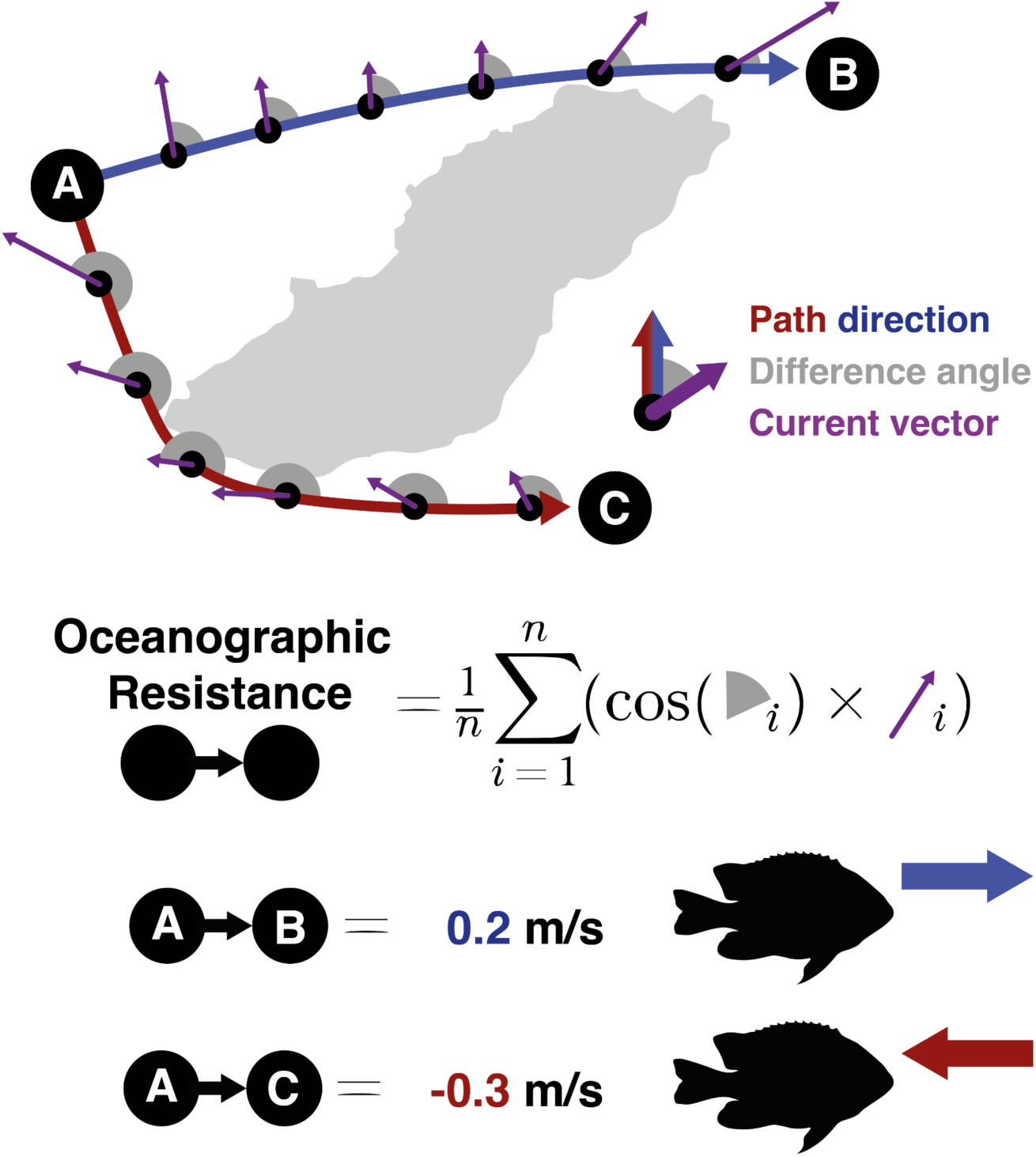
A visual definition of oceanographic resistance, a metric to describe the current system vectors encountered between two points. Simplified examples of calculating oceanographic resistance from point A to B and point A to C around an example island (San Cristóbal, Galápagos) are shown. The angle of difference between the path direction and current direction is calculated at equally spaced points along the shortest ocean path between two points. At each point the cosine of the angles is multiplied by the current magnitude, oceanographic resistance is the mean of the resultant value across the entire path (measured in mean metres per second). Oceanographic resistance is negative for currents against the travel direction on average and positive for currents in the same direction of travel on average.

The mean oceanographic resistance was 0.00 ± 0.14 (s.d.) with a maximum value of 0.293 and a minimum value of -0.292. A comparison of Lagrangian minimum drift days and the oceanographic resistance values for journeys for which minimum drift could be calculated revealed a weak correlation (linear regression, R^2^ = 0.086, p < 0.001), driven entirely by the highest values of drift days occurring with the lowest oceanographic resistance (see Fig. S3). There was no relationship (linear regression, p > 0.05) between oceanographic resistance and minimum drift days excluding the lowest values of oceanographic resistance (< -0.1 as shown in Fig. S3). Collectively, this indicates that, compared to Lagrangian particle tracking metrics, oceanographic resistance captures novel and non-overlapping characteristics of the current systems integrated across a journey between two sites.

### Fine-scale ocean currents influence local fish biodiversity

We found a positive relationship (distance-decay) between eDNA-measured site dissimilarity and geographic distance between pairs of sites (Fig. 3). There was a significant relationship between site dissimilarity and geographic distance, temperature difference and oceanographic resistance (Maximum-likelihood population-effects generalised least squares regression, p < 0.001 for all variables, Table S3, Figure S4).

**Fig. 3.**
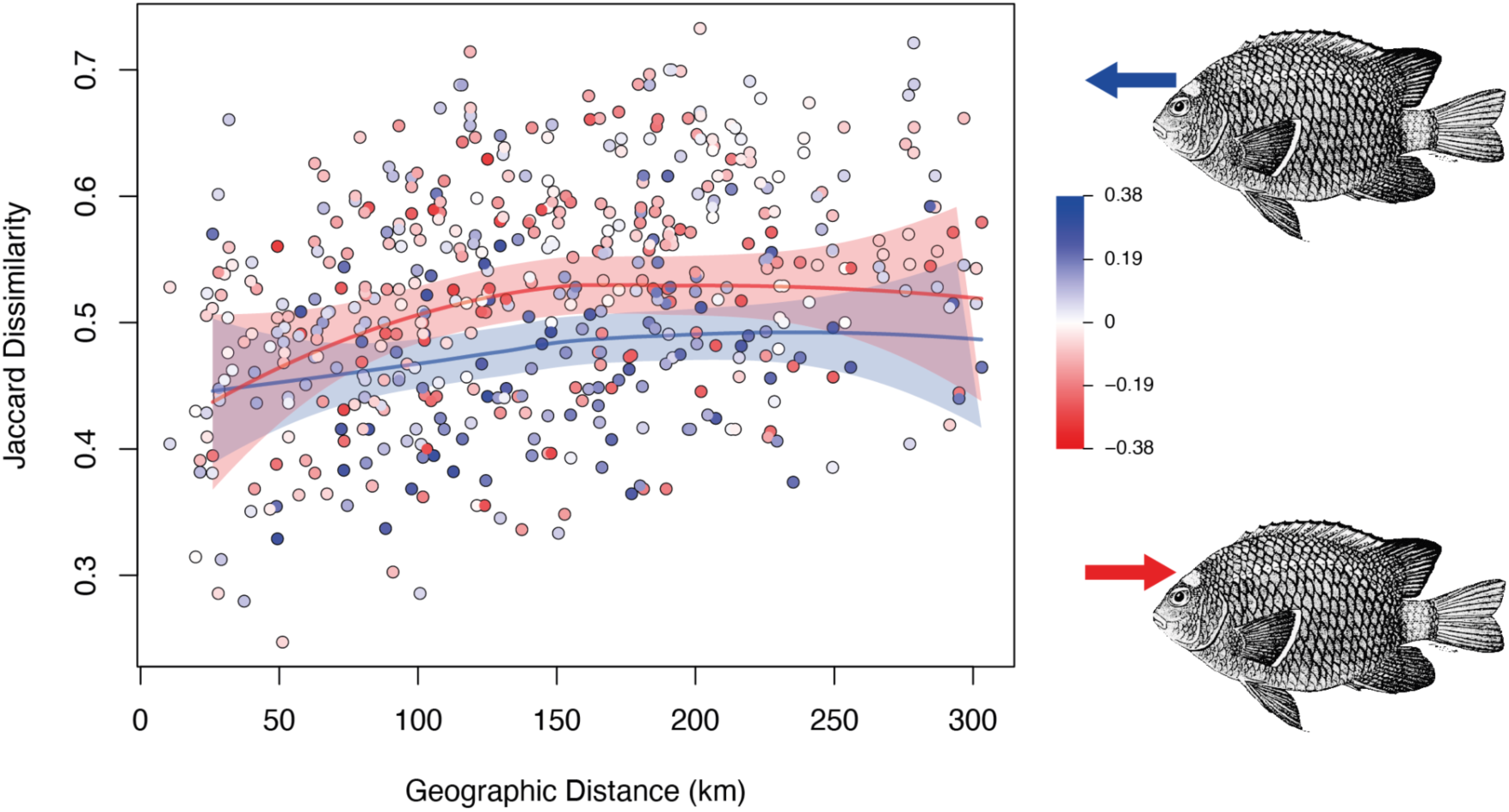
Modified asymmetric Jaccard dissimilarity for each pair of sites, displayed against geographic distance measured in km. Each point is coloured according to the oceanographic resistance between pairs of sites; point colour indicates oceanographic resistance with scale from -0.38 m s^-1^ to 0.38 m s^-1^ shown on the right legend. Loess smoothed fit lines for data below the 20^th^ percentile and above the 80^th^ percentile of oceanographic resistance are shown as red and blue lines respectively, with shading indicating the 95% confidence interval of the fit. Damselfish (Stegastes beebei) illustrations ^42^ on the right denote the direction of average current flow for highly positive (blue) and highly negative (red) oceanographic resistance.

Across the distance-decay relationship when oceanographic resistance was negative, sites were on average more dissimilar than in cases where oceanographic resistance was positive (Fig. 3). A similar trend was observed with temperature data, with greater dissimilarity between sites on average when temperature change between sites was positive (Fig. S5).

A maximum-likelihood population-effects (MLPE) generalised least squares regression with the smaller number of observations (243 journeys between sites) provided by the Lagrangian particle tracking found a significant effect of geographic distance and temperature difference on eDNA-measured site dissimilarly (p < 0.05), but found no significant effect of oceanographic resistance or Lagrangian minimum drift days (p > 0.05); full model outputs are provided in Table S4 and Fig. S6. In agreement with MLPE models, a Mantel test found a significant (r = 0.33, p < 0.001) correlation between eDNA-measured site dissimilarity and geographic distance. However, no significant correlation between eDNA-measured site dissimilarity and oceanographic resistance or temperature (p > 0.05) was detected. A Mantel test between eDNA-measured site dissimilarly and Lagrangian minimum drift days was not performed due to missing observations where particles did not drift to the journey site destination.

In order to evaluate if the number of fish detected (ASV richness) was correlated with the spread of eDNA through time by ocean circulation, Lagrangian particle release experiments were conducted from each site run back in time for 72 hours, based on previous studies of marine eDNA decay ^40,41^, to estimate possible eDNA contributions for each sampling event. No significant (p > 0.05) relationship was found between ASV richness and all four calculated metrics of oceanographic spread (mean distance of particle centroid from sampling site, mean spread of particles around centroid, surface area occupied by particles, mean of individual particle distance from sampling area) (Fig. S7).

## Discussion

We found heterogenous fish community structure across the Galápagos islands, with eDNA metabarcoding-measured beta diversity patterns generally agreeing with previously described bioregions ^30^. Remarkably, we found not only that variation in fish communities could be explained with the submesoscale flow data generated by our ocean circulation model, but that the magnitude of the effect of currents was similar to temperature, a well-known determinant of marine biodiversity ^17,43^. These findings relied on the development of a new metric, oceanographic resistance - which describes unique variability in ocean currents not incorporated by Lagrangian approaches. Overall, these results advance our understanding of the determinants of fish biodiversity from a unique archipelago, and provide a novel metric to investigate the role of submesoscale currents on ecosystems across the globe.

Given that previous work has described fish bioregionalization across the archipelago ^30^, it is unsurprising that eDNA metabarcoding provides similar evidence for fish biogeography. Consequently, these patterns underline the unique nature of the Galápagos, but more broadly show unusually clear differences in fish communities across short (<200 km) geographic distances. Many eDNA surveys have found biogeographic regionalization, particularly changes in community structure (beta diversity) in marine ecosystems ^43–47^. However, studies in other regions have shown that marine fish communities can also have homogenous community structure, even across large (>1000 km) distances ^48–50^. Collectively these previous investigations suggest that homogenous biogeographic structure should be our null hypothesis for communities of highly mobile marine organisms at local regions.

An important novel piece of biogeographic evidence in our study is the unexpected grouping of Roca Redonda in the Northern bioregion (Fig. 2). This should prompt further research to investigate the, here un-sampled, Far-Northern Islands (Darwin & Wolf), which may have unanticipated biodiversity. This could potentially require a change in bioregion designation and thus management strategy. Given the limited sampling of the Elizabeth bioregion, further work is required to understand if, and how, fish communities in this region differ from the surrounding Western bioregion.

Bioregionalisation is likely a consequence of the unique oceanographic conditions around the Galápagos islands ^30–32^. We found that Lagrangian particle modelling did not explain a significant proportion of either alpha or beta diversity in the fish communities of the Galápagos. This stands in contrast to other work in which particle tracking explained variation in whole-community dissimilarity in Mediterranean ^10^ and Pacific ^9^ coastal communities. Fish communities are variably affected by currents. Many have planktonic dispersal stages ^14,51^, others have tightly defined home ranges, and some are nektonic - travelling vast distances to forage ^17^. In sum, fish communities do not act in a similar way to Lagrangian particles, motivating our development of a new metric to describe the current systems across a seascape journey. This metric enabled us to describe current variability across journeys for which Lagrangian metrics could not be calculated. However, just as fish do not act like passive particles, they do not act as perfect agents moving through the shortest seaward distance as described in the derivation of oceanographic resistance. Future work should explore the likelihood of various paths through the ocean that could be used to calculate oceanographic resistance, to improve the estimation of the resistance experienced across the seascape. Despite utility in explaining differences between sites, our metric ignores many variables that may have an important influence on current systems, for example, depth, variability across time, and variation in the mean current experienced across the journey.

Our analysis combining novel oceanographic modelling and eDNA metabarcoding data could only explain a small proportion of the total variation among sites using distance and temperature data (Fig. 2 and Fig. S4). Studies evaluating the explanatory power of a set of environmental and spatial predictors typically only describe a small fraction of the total beta diversity in marine communities ^43,52,53^. These findings are also reflected in meta-analyses across ecosystems, with much of the measured variation in communities remaining unexplained ^54–56^. Metacommunity theory predicts that ecological drift (i.e. stochastic demographic changes in species composition) is likely to occur under both neutral and selective population dynamics ^57^, and thus some variation in community composition will always be unexplained by environmental and spatial predictors. Furthermore, there is frequently a compromise between surveying across space and through time to capture community dynamics, with even the most comprehensive ocean surveys showing only a snapshot of temporal dynamics ^5^. Moreover, while eDNA methods for fish detection are well developed ^24^, it is likely that future methodological improvements can increase the detection of some species.

Our MLPE analyses indicated that geographic distance, temperature and oceanographic resistance were important explanatory variables describing patterns in beta diversity in Galápagos fishes. Moreover, we demonstrate that the increase of power gleaned by oceanographic resistance, in comparison to Lagrangian particle tracking, allowed us to find an effect of current on community composition. Given ongoing controversy concerning the power and appropriateness of Mantel tests to test for association between matrices^58–60^, it is unsurprising that these analyses provided support for only the strongest effects in the dataset, namely a distance-decay effect. Distance-decay relationships (changing biotic composition across space) are well studied in both marine and terrestrial ecosystems; thus an effect of geographic distance was expected ^8^. Similarly, temperature has been shown to be a key variable structuring communities of both fish ^61^ and other marine organisms ^5,13^. Work exploring the effect of current systems on marine biodiversity has either combined geographic and current-based distance ^9^, or been limited to benthic marine organisms ^10^. In line with our findings, these studies do find an important role for fine-scale ocean currents in structuring marine communities. Fish have both nektonic and planktonic phases to their life cycle, with planktonic eggs and larvae developing into nektonic adults^51^, thus our expectation should be a reduced or equal effect of currents on community difference compared to studies of less mobile organisms. We showed that oceanographic resistance contains unique explanatory information, demonstrating that the direction and magnitude of currents connecting sites can influence the composition of fish communities. Given the significant effect of human-induced climate change on ocean currents and mixing ^62^, work is urgently needed to assess how currents on such fine scales will affect biodiversity patterns in other taxa and ecosystems.

## Methods

### Study area

The Galápagos Archipelago is made up of 13 major islands ranging in isolation from 6-46 km from their nearest neighbour, lying in the Eastern Tropical Pacific Ocean, approximately 900 km west from mainland Ecuador. Previous diver-based rocky reef surveys of fish and macroinvertebrates in shallow coastal waters around the islands ^30^ revealed a marked bioregional signal across the archipelago. The five main bioregions identified were a far northern region around the remote islands of Darwin and Wolf; a warm northern region encompassing the islands of Pinta, Genovesa and Marchena; a mixed region around the central islands; a cool western region around Fernandina and western Isabela; and a region consisting of the channel separating Isabela and Fernandina, including Elizabeth Bay, sufficiently distinct from the western bioregion to merit status as an independent bioregion (see Fig S2).

### Sample collection

We collected seawater samples from 23 sites across the southern and central Galápagos Islands (Fig.1a) during September 2018 (see Table S2 for details). At each sampling point 1 L of seawater was collected from 30 cm below the surface with a Kemmerer water sampler (cleaned with 5% bleach solution each time before use) and filtered through a 0.22 μm polyethersulfone Sterivex filter (Merck Millipore, Massachusetts USA) using a sterile 50 ml luer lock syringe. Additionally, 2 L of seawater were collected at the maximum depth of each site (ranging from 3.5 to 30.5 metres) and filtered using the same method, resulting in a total of 3 L of water per site. As metazoan diversity detected by eDNA varies across depth ^63^, this approach aimed to characterise total fish diversity at the site. To minimise contamination among samples, reused equipment was cleaned with 5% bleach solution. A negative control sample consisting of filtered distilled water sample was taken for each deployment. We added 2 ml of ATL Buffer (Qiagen) to each Sterivex filter to preserve eDNA and stored them at room temperature until further processing.

### DNA extraction and library preparation

We used the dedicated low-DNA laboratory at the National Oceanography Centre, Southampton (United Kingdom) to conduct the DNA extraction. This laboratory was separated from facilities where PCR was performed. No high copy DNA template, cultures or PCR products were permitted in this laboratory. All laboratory surfaces, reused equipment and reagent packaging were thoroughly cleaned with 5% bleach solution. DNA was extracted following the SX^CAPSULE^ method from ^64^, with sample identifiers blinded before extraction to avoid unintentional human bias. The final DNA elution was performed with 200ml DNase free water and an additional re-elution was performed with the eluate. Marine eDNA samples can contain PCR inhibitors, which have a negative effect on species detection sensitivity ^65^. We therefore tested for inhibition using a Primer Design Internal Positive Control qPCR Kit (Southampton, United Kingdom) and Primer Design Precision Plus Mastermix with 20 µl reactions containing 4 µl of eDNA for each sample under the manufacturer recommended conditions. Inhibited samples were expected to have an increase in Ct (cycle threshold) of >1 compared to the unspiked reaction. As inhibition was detected in a fraction of sites, all samples were treated using the Zymo OneStep PCR Inhibition Removal Kit (Zymo Research, Irvine, USA) following the manufacturer recommended protocol.

We used metabarcoding primers that targeted a variably sized (163-185 bp) fragment of the mitochondrial 12S region ^66^. These primers consist of two sets, one targeting teleost fish, and a second set targeting elasmobranchs (sharks and rays). The entire metabarcoding PCR and library build was performed independently for these two primer sets. Metabarcoding libraries were constructed using a 2-step method where an initial PCR incorporates an adaptor sequence onto the 5’ end of the primers that serves as the target for a second PCR that incorporates index sequences for demultiplexing and Illumina sequencing adaptors (following Holman et al. 2021). For each set of primers PCR reactions were conducted in 20 µl volumes consisting of 10 µl AmpliTaq Gold 360 mastermix (Agilent Biosystems, Waltham, USA), 1.6 µl of primers (5 µM per primer) and 2 µl of undiluted cleaned eDNA template. The reaction proceeded with an initial hold at 95°C for 10 minutes followed by twenty cycles of 95°C for 30 seconds, 59°C for 30 seconds and 72°C for 60 seconds, a terminal hold at 72°C was conducted for 10 minutes. As the number of PCR replicates per sample increases the diversity detected ^67^, we conducted eight independent replicate reactions per water sample that were then pooled for bead cleaning and indexing. These pools were cleaned using Beckman Coulter (Brea, USA) AMPure XP beads, for each 160 µl pool (eight 20 µl reactions), 128 µl of beads were added and the manufacturer recommended protocol was followed with a final elution of DNA into 20 µl of 10mM Tris-HCl buffer (pH 8.5). The second PCR was conducted in 20 µl volumes consisting of 10 µl AmpliTaq Gold 360 mastermix (Agilent Biosystems, Waltham, USA), 1.0 µl of primers (10µM per primer), and 5µl of bead-cleaned first PCR product. The reaction proceeded with an initial hold at 95°C for 10 minutes followed by fifteen cycles of 95°C for 30 seconds, 55°C for 30 seconds and 72°C for 60 seconds, a terminal hold at 72°C was conducted for 10 minutes. The product was then bead cleaned as above with 16 µl of beads in each 20µl reaction. Libraries were then individually quantified using the New England Biolabs (Ipswich, USA) NEBNext Library Quant Kit and pooled at equal molarity into two libraries, one for each initial primer set. These two libraries were diluted to 4 nM, spiked with 5% PhiX for diversity and sequenced in two independent runs of a Illumina (San Diego, USA) MiSeq instrument using a V3 2x300 bp kit. Negative controls from sampling, DNA extraction and PCR one and two blanks (RT-PCR grade water) were all amplified, pooled and sequenced along with experimental samples.

### Bioinformatic analyses

Raw sequences were demultiplexed using the GenerateFastQ (v.2.0.0.9) module on the MiSeq control software (v.3.0.0.105) under default settings. Primers were then trimmed from both paired reads, ensuring both the forward and reverse primer was present in each read pair using Cutadapt ^68^ (v.3.2). As the sequencing length covered the entire target amplicon, the reverse complement of each primer was also trimmed using Cutadapt from the 3’-end of each read pair. Following primer trimming reads were denoised into ASVs using the DADA2 pipeline (v1.16.0) ^69^ in R ^70^ (v.4.0.3) under default parameters unless detailed below. The *filterAndTrim* function was conducted using a maxEE value of 1 and truncLen value of 120 bp for both read pairs. After the generation of an ASV table the data was curated by running lulu (v.0.1.0) ^71^ under the default parameters. Each independently sequenced dataset was then cleaned separately as follows using R. Positive observations were discarded if they had fewer raw reads than the sum of all reads found in the negative control samples for each ASV, or if they had fewer than three reads. ASV by sample tables were then transformed into proportional abundance per sample and data from identical ASVs was merged using the collapseNoMismatch function in DADA2. It is commonplace to multiplex the two MiFish primer sets (elasmobranch and teleost) during PCR and treat them as a single marker ^66,67^, because they differ by only three nucleotides across the forward and reverse primers, they amplify many species in common. However, we instead chose to increase the sequencing output per sample and conservatively merged these primer sets bioinformatically as above. ASVs were then queried against the entire NCBI *nt* database (updated on 01-02-2021) using a BLAST+ (v.2.11.0) nucleotide search with *-num_alignments* set to 200. These alignments were then filtered using the R script ParseTaxonomy (DOI:10.5281/zenodo.4564075) that was previously developed ^43^ to generate lowest-common-ancestor assignments in the case of multiple matches. Initial analyses revealed some errors in the assignments, likely due to missing taxa in databases, so all ASV assignments were curated using the online NCBI blastn portal (accessed March-August 2022). Erroneous sequences were identified as having no match to any nucleotide or protein (using a blastx search) 12S sequence, all such sequences were discarded. ASVs matching domestic animals (cow, dog, chicken etc.) or human DNA were removed from the main dataset. Finally, ASVs with an unambiguous species assignment (>99% sequence similarity across the whole sequence, matches to other species in the genus >1% sequence similarity from the proposed assignment) were merged.

### Oceanographic analyses

Particle tracking simulations were conducted using a realistic ocean general circulation (OGCM) model constructed using MITgcm^72^ with a 1 km horizontal resolution covering an area encompassing the Galápagos Marine Reserve. This 1 km model has 630 grid points in the zonal direction, 768 grid points in the meridional direction, a grid origin at 3.1°S 94.1°W, and four open boundaries. The 1 km model is embedded within a larger regional OGCM with a model grid resolution of 4 km in the horizontal between ± 5° latitude, stretching out to ∼12 km in latitude at the model boundaries. Both 1 km and 4 km models were constructed using bathymetry from the General Bathymetric Chart of the Oceans ^73^ grid and a vertical grid comprising 75 depth levels, with vertical resolution varying with depth from 5 m over the first 50 m, 9.8 m to 164 m depth, and 13.7 m to 315 m depth, and a maximum cell height of 556 m below 3000 m. Boundary forcing and initial conditions for the 1 km model were taken from the 4 km model. The 4 km model was run with three completely open boundaries (North, South and West), using periodic boundary forcing for temperature, salinity and velocity fields and a 15-grid box thick sponge layer for velocity. Initial conditions and monthly boundary forcing were taken from the Mercator Ocean reference model^74^, a global ocean model based on 1/12 (0.083) degree NEMO^75^. Atmospheric forcing, wind stress and evaporation and precipitation for both models were taken from the ERA-Interim^76^ reanalysis at a 3-hour temporal resolution for all fields, and radiation (shortwave and longwave) forcing from Modern-Era Retrospective analysis for Research and Applications (v.2) (MERRA2^77^) at hourly temporal resolution.

Particle tracing experiments were performed in the 1 km model using TRACMASS^78^ to establish the likely origin of waters at the sample sites. Particles were seeded into the model at a concentration of 1 particle per 3x10^3^ m^3^ volume, covering a horizontal area of ∼4 km^2^ around the sample site, from the surface to 20 m depth. Particles were released five times at each sample site (2 days before to 2 days after) sampling. Particles were then tracked backwards-in-time through the model flow field for 3 days. The final positions of particles from all releases were aggregated and normalised (as a fraction, where one is the sum of all particles released) and a spatial distribution for likely sample site water origin estimated. Four parameters describing the spread of the particles 72 hours before sampling were calculated: the direct line distance between the average latitude and longitude of the points from the release point, the mean distance of the particles from the mean latitude and longitude of the points, the surface area occupied by grid squares with greater than 0.01% of released particles, and the average of the individual particle direct line distances from the release point (see example particle drift in Fig. S8).

Drift times between sites were estimated by particle tracing in the 1 km model. Particles were released as described above at each sample site and tracked backwards through the model flow field for 60 days. The shortest time for particles from each of the other sample locations to arrive at the source sample site was recorded (i.e. days for the first particle from the source sample site to reach the other sample locations) along with the fraction of the released particles arriving at the other sample locations during the 60 day trace.

### Calculating oceanographic resistance

The General Bathymetric Grid of the Oceans 2022 grid ^73^ was subset around the Galápagos Islands using the sf package (v.1.0.9.) ^79^ in R (v.4.2.2). This dataset contains seawater depth and coastline information at a resolution of 15 arcseconds. For each possible journey from every site to every other site, the shortest path avoiding land masses was calculated using the *shortestPath* function in the *gdistance* R package (v.1.6) ^80^. These data are henceforth referred to as geographic distance. To estimate the overall water resistance experienced by an agent travelling along the shortest path in the study area between two sites, taking into account ocean currents, we devised a metric that we refer to as oceanographic resistance (see Fig. 2), which differs to the previously described ^81^ ‘derived oceanographic resistance’ as it calculates the oceanographic forces across a journey and not the ‘derived’ values of population connectivity as Thomas et al. 2015 ^81^.

For each site-to-site geographic distance, a point was extracted from along the path every 1 km using the *spsample* function from the *sp* R package (v.2.1-4) ^82^. Northings were extracted from the 1 km model as mean monthly northward velocity (m s^-1^, positive going north) and Eastings as mean monthly eastward velocity (m s^-1^, positive going east) from the 1 km model for 2018. From these Northing and Easting values, the resultant vector was calculated and represented by magnitude and azimuth degrees. The azimuth of the oceanographic current for each extracted point was then compared to the azimuth between the extracted point and the subsequent point along the path. The resultant angle indicates the difference between the direction of travel and the direction of the current, with for example, zero degrees indicating that the current and direction of travel are identical and 180 degrees indicating that the current and direction of travel are opposite. This comparison angle was then transformed using a cosine function to give a value of 1 and -1, respectively for the previous examples. The oceanographic resistance at the extracted point was calculated by multiplying the result of the cosine function by the magnitude of the current at the point. Finally, the oceanographic resistance for a given path was calculated as the mean of the oceanographic resistance of all selected points on the path between the start and end points. Oceanographic resistance is a mean value of a series of transformed vectors measured in m s^-1^, and as such is a scalar measured in m s^-1^.

### Ecological analyses

All analyses were conducted in R (v.4.2.2) unless otherwise stated. Differences in mean ASV richness between bioregions were evaluated using a one-way ANOVA. Beta dissimilarity between sites was visualised with non-metric multidimensional scaling (nMDS) using a Jaccard dissimilarity, an index appropriate for testing biogeographical patterns ^83^, implemented with the metaMDS function from vegan (v.2.6-4) ^84^. All beta diversity analyses were conducted on averaged values among the three replicates per site. Subsequent statistical tests on bioregions followed the designation of ^30^ with one site (Roca Redonda) in a previously unsurveyed region placed in a bioregion according to the results of the nMDS. Differences in within-bioregion multivariate dispersion were evaluated using the PERMDISP ^85^ procedure implemented in *betadisper* function from vegan, with post hoc testing of pairwise differences tested using the *TukeyHSD* function. Statistically significant differences between bioregions were evaluated using a PERMANOVA ^86^ on Jaccard dissimilarities implemented using the *adonis2* function in vegan. Pairwise PERMANOVA comparisons between bioregions were implemented using the *adonis.pair* function in the EcolUtils package (v.0.1) ^87^.

In contrast to oceanographic resistance, Jaccard dissimilarity is symmetric considering the order of sites. For example, for a pair of sites, the oceanographic resistance defined above may differ depending on the direction of travel from site A to site B, while the Jaccard dissimilarity measures the difference between sites symmetrically. In order to test for an effect of oceanographic resistance on beta diversity a modified asymmetric Jaccard dissimilarity was implemented such that the order of the two sites supplied to the function (i.e. Site A to Site B / Site B to Site A) changed the output as below.

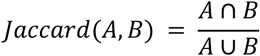

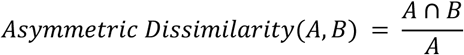

This modified dissimilarity measure can be interpreted as the dissimilarity between site A and site B considering only species present in site A. In other words, species not found in site A that are present in site B do not contribute to the dissimilarity index. Additionally to evaluate the comparative effect of oceanographic resistance to other marine conditions we extracted the average October 2018 mean sea temperature from the top 20 metres of the model output for each site, transforming these values into temperature differences between sites to compare against distance matrices.

A maximum-likelihood population-effects^88^ generalised least squares model was performed in R using the *gls* function from the *nmle* package^89^ (v.3.1-166) as follows. Asymmetric dissimilarity was used as the dependent variable against geographic distance with an additive effect of oceanographic resistance and temperature difference with the “REML” method and a correlation structure provided with the *corMLPE* function from the *corMLPE* package (v.0.0.3) with the form ‘∼ Start + End’ used to account for the non-independence of observations from and to the same site.

A second generalised least squares model was constructed with the same form but with minimum Lagrangian particle drift days added as an additional additive variable to examine the effect of variables with the loss of power due to the required data subsetting as Lagrangian descriptions of all journeys are not possible as outlined in the results.

Mantel tests were performed with a Pearson coefficient test statistic with 9999 permutations to test for significance. This test was implemented using custom R code presented alongside this manuscript to test and permute across the entire matrix (upper and lower sections) excluding the diagonal.

To test for possible correlations between ASV richness and particle spread, least-square regression models were implemented using the function *lm* with each of the particle spread statistics described above.

All authors thank the Galápagos National Park Directorate (GNPD), Universidad San Francisco de Quito (USFQ), and Galapagos Science Center (GSC) for logistic support for fieldwork, and the Ministry of Environment, Ecuador for granting a collection permit (MAE-DNB-CM-2016-0041). The work received support from a UK Royal Society Grant (CH160019) led by ANG. LH was supported by the Natural Environmental Research Council UK (grant number NE/L002531/1) and the European Research Council (ERC) under the European Union’s Horizon 2020 Research and Innovation Programme (grant agreement no. 856488). MR acknowledges support from the MARGECH research grant (PID2020-118550RB) of the Spanish Ministry of Science and Innovation (MCIN/AEI/10.13039/501100011033). MR and LH acknowledge support from the BlueDNA research grant (PID2023-146307OB) funded by the Spanish Ministry of Science, Innovation, and Universities (MICIU/AEI/10.13039/501100011033) and by ERDF/EU. The authors thank Julián Regalado Pérez for discussions on oceanographic resistance.

## Author Contributions

ANG initiated the project and secured funding, supported by DP, AH and MR. LH, DP and MR designed the field sampling and samples processing strategy. DP conducted the fieldwork. LH conducted the laboratory work, bioinformatics, ecological analyses and wrote the initial manuscript. AF and ANG conducted oceanographic modelling analyses. All authors substantially contributed to further manuscript drafts and provided final approval for publication.

## Data & Code availability

The raw Illumina sequencing data are available from the European Nucleotide Archive under study accession number PRJEB55415. All other metadata, intermediate data and scripts are permanently archived at DOI: 10.5281/zenodo.10593433.

