## Supplementary Information for "Fine-scale oceanographic processes shape marine biodiversity patterns in the Galápagos Islands"

### Supplementary Text 1 - Sequencing metrics and controls

The universal and elasmobranch Illumina sequencing runs produced 9,276,041 and 14,801,645 raw paired end reads respectively that could be assigned to a sample index. The run using universal primers produced an average of 132,031.3 ± 60,736.9 (s.d.) raw reads per non-control sample with 125,825.2 ± 57,427.0 (s.d.) reads remaining after low quality and chimeric reads were excluded. The run using elasmobranch primers produced an average of 212,978.1 ± 1,063,729.3 (s.d.) raw reads per non-control sample with 190,523.5 ± 933,726.9 (s.d.) reads remaining after low quality and chimeric reads were excluded, the high variance in raw read output was due to three samples receiving a larger than expected proportion of reads (7,675,465; 1,377,703; 895,459) due to a pooling error. Excluding these samples from the above calculations resulted in 53,120.2 ± 20,338.5 (s.d.) raw and 48,446.9 ± 18,638.9 (s.d.) filtered reads per sample. In total 23,111 reads were found in the negative control samples across both runs with 15 and 42 ASVs in the universal and elasmobranch datasets respectively. The majority (81.8%) of these reads were from 12 ASVs assigned to homo sapiens, and 348 reads were from 12 ASVs assigned to fish species. After cleaning and filtering 485 and 563 ASVs were retained for the universal and elasmobranch datasets respectively. The merging of these datasets and removal of human, domestic and all other ASVs not assigned to fish (teleost and elasmobranch) species resulted in a dataset of a total of 551 ASVs, which was then used for all subsequent analyses. Taxonomic assignment indicated all ASVs were from elasmobranch or teleost fish, with 66 ASVs assigned to species, 216 to Genus, 167 to Family and 99 above Family.

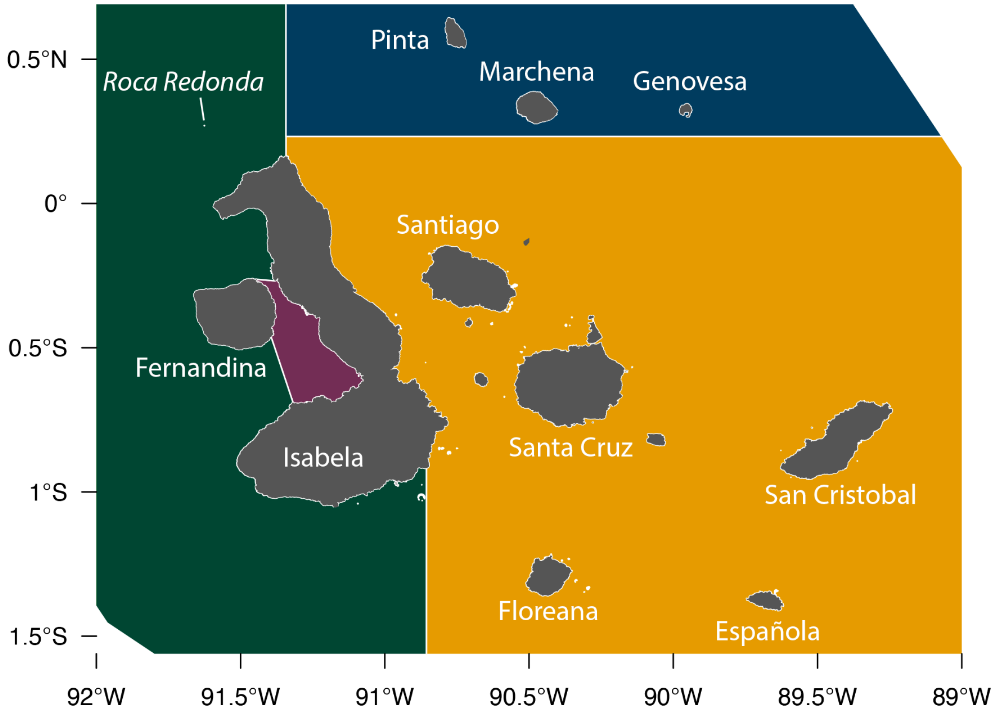

**Fig. S1.** Labeled map of the Galápagos islands. Bioregions are shown by the color with the western bioregion in green, the Elizabeth bioregion in purple, the Northern bioregion in blue and and Central South-eastern bioregion in orange.

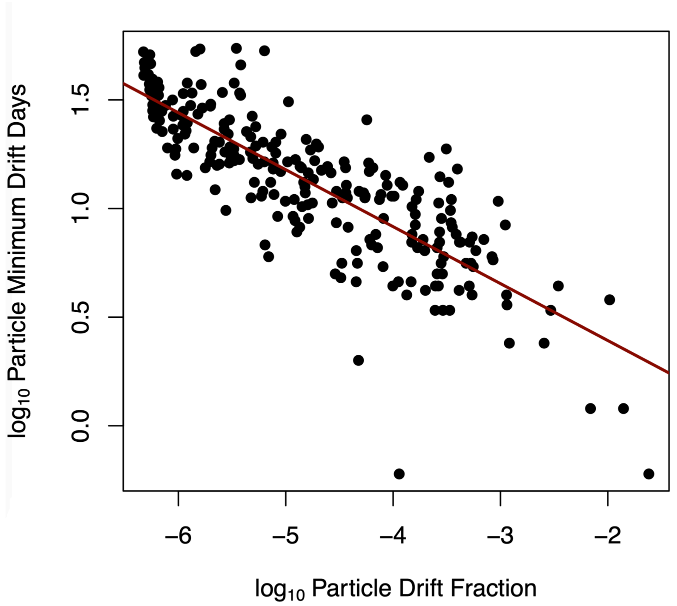

**Fig. S2.** Relationship between drift fraction and drift days. The scatter plot shows the log-transformed values of drift fraction (measured as a fraction) and drift days (measured in days). The red line represents the best-fit regression line from the linear model indicating a strong negative correlation between the two variables.

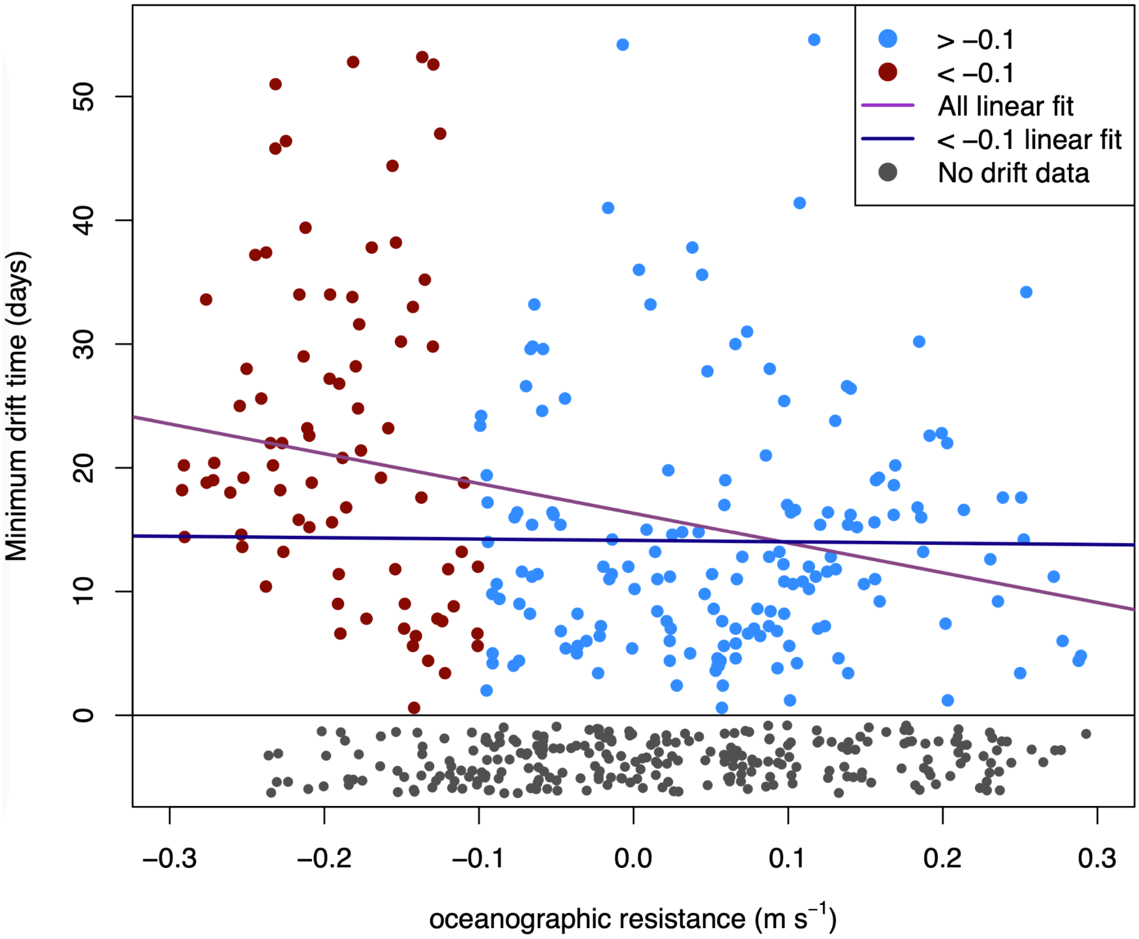

**Fig. S3.** Relationship between oceanographic resistance (m s$^{-1}$) and minimum drift time measured in days. Blue points represent resistance values > -0.1, and red points represent values < -0.1. The purple line shows the linear regression fit across all data, while the blue line shows the fit for values > -0.1. Grey points indicate cases where drift time data could not be generated, jittered vertically for clarity.

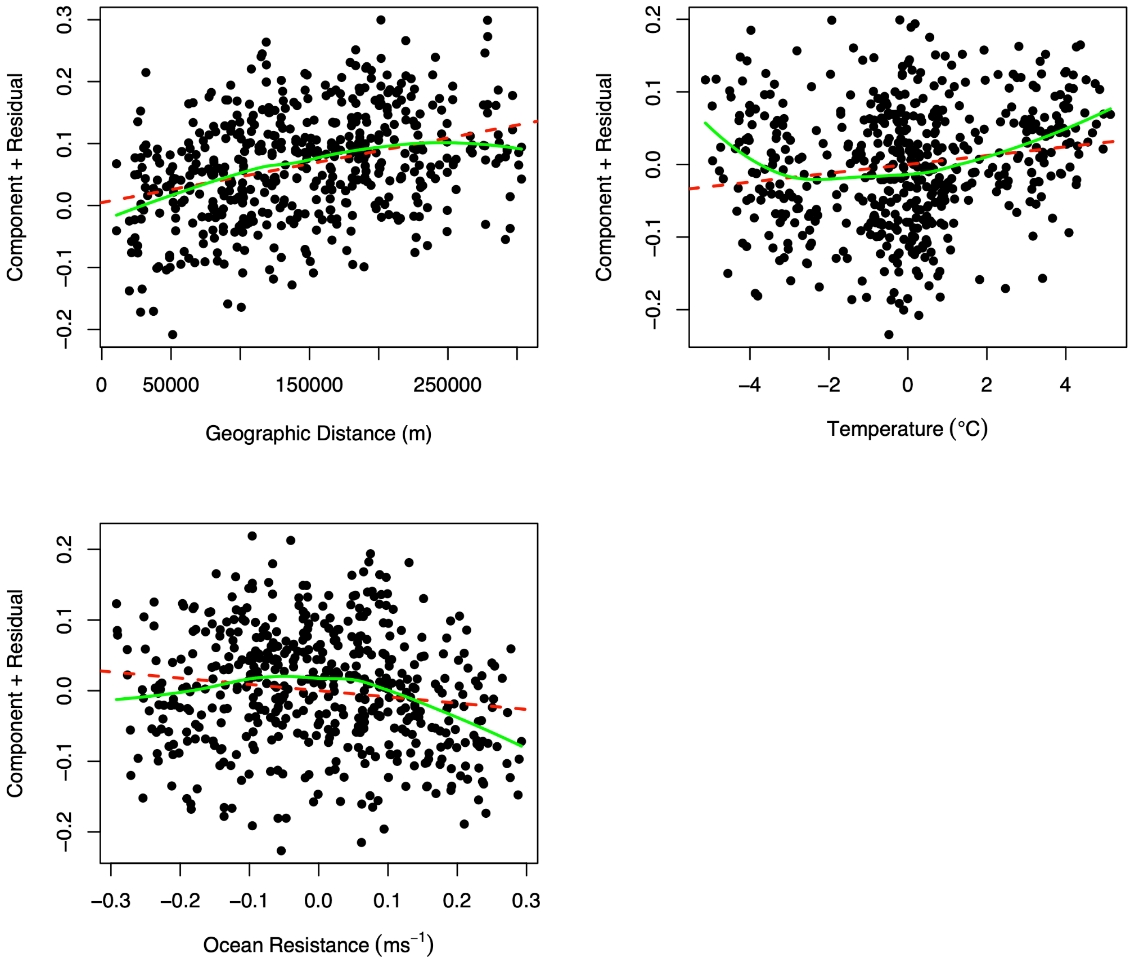

**Fig. S4.** Component + residual plots (partial residual) for a multiple linear regression model predicting eDNA-derived asymmetric Jaccard dissimilarities from (top left) geographic distance (m), (top right) temperature difference (°C), (bottom left) oceanographic resistance ($ms^{-1}$). The fit for each model variable is shown by red dashed line, a loess fit of the data is shown by a solid green line.

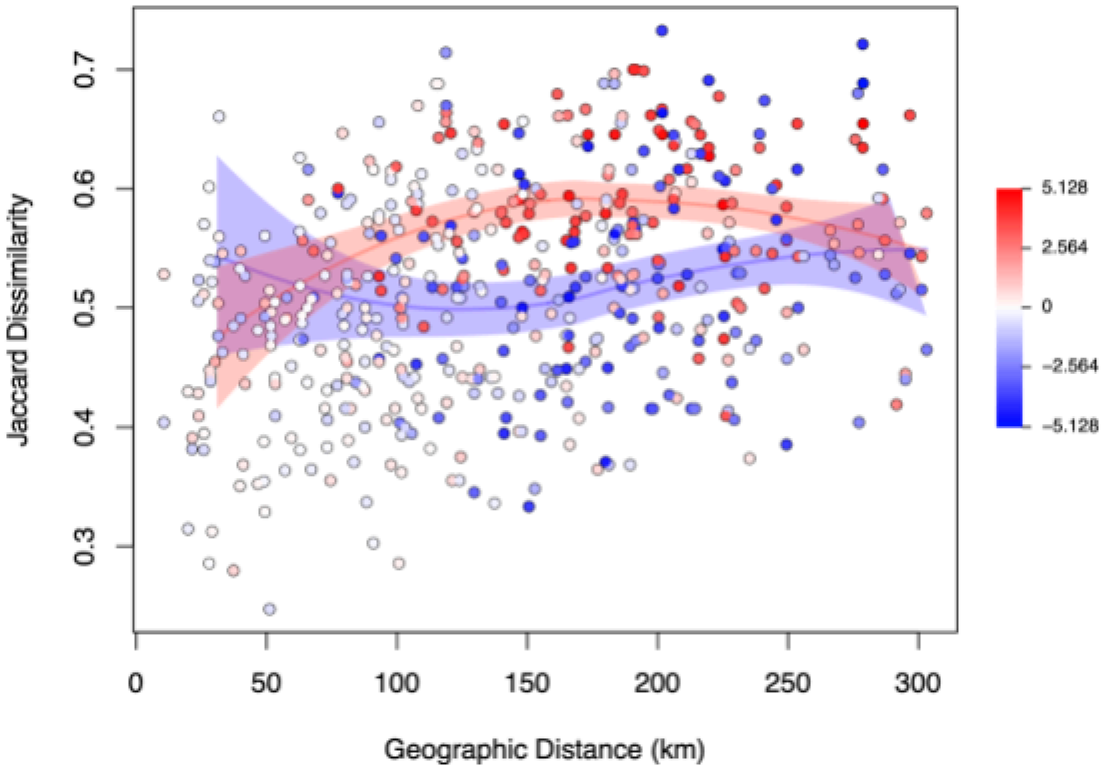

**Fig. S5.** Modified asymmetric Jaccard dissimilarity for each pair of sites, displayed against geographic distance measured in km. Each point is colored according to the temperature difference between pairs of sites; point color indicates temperature difference with scale shown on the left, measured in °C. Loess smoothed fit lines for data below the 20th percentile and above the 80th percentile of oceanographic resistance are shown as red and blue lines respectively, with shading indicating the 95% confidence interval of the fit.

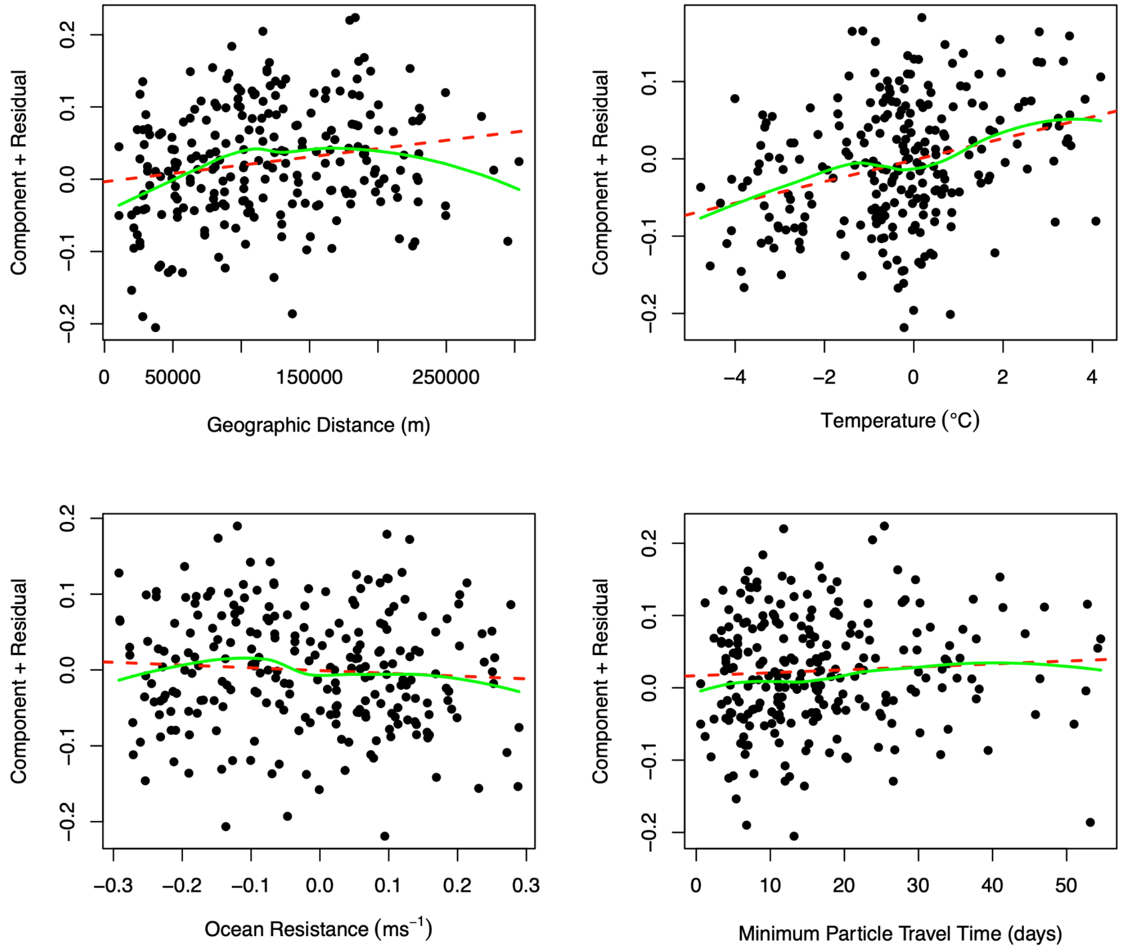

**Fig. S6.** Component + residual plots (partial residual) for a generalized least squares linear regression model predicting eDNA-derived asymmetric Jaccard dissimilarities from (top left) geographic distance (m), (top right) temperature difference (°C), (bottom left) oceanographic resistance ($ms^{-1}$) and (bottom right) Lagrangian minimum drift days. The fit for each model variable is shown by red dashed line, a loess fit of the data is shown by a solid green line.

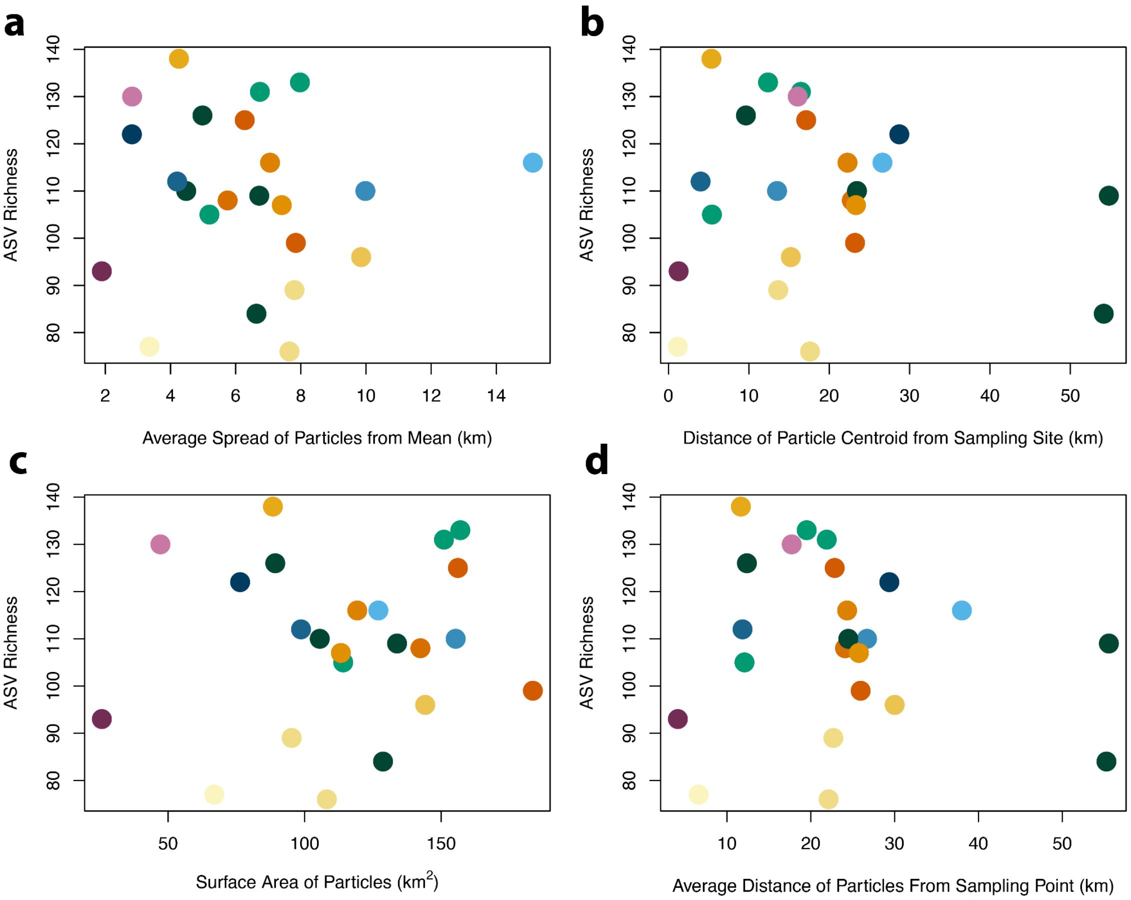

**Fig. S7.** Measures of particle spread from the collection site 72 hours before eDNA sampling compared to ASV richness at each site. Colors from points indicate sites matching those in Fig. 1 from the main manuscript with green points from the western bioregion, blue from the northern bioregion, purple from the Elizabeth bioregion and orange/yellow from the central south-eastern bioregion. a) The direct line distance between the average latitude and longitude of the points from the release point b) the mean distance of the particles from the mean latitude and longitude of the points, c) the surface area occupied by grid squares with greater than 0.01% of released particles d) the average of the individual particle direct line distances from the release point.

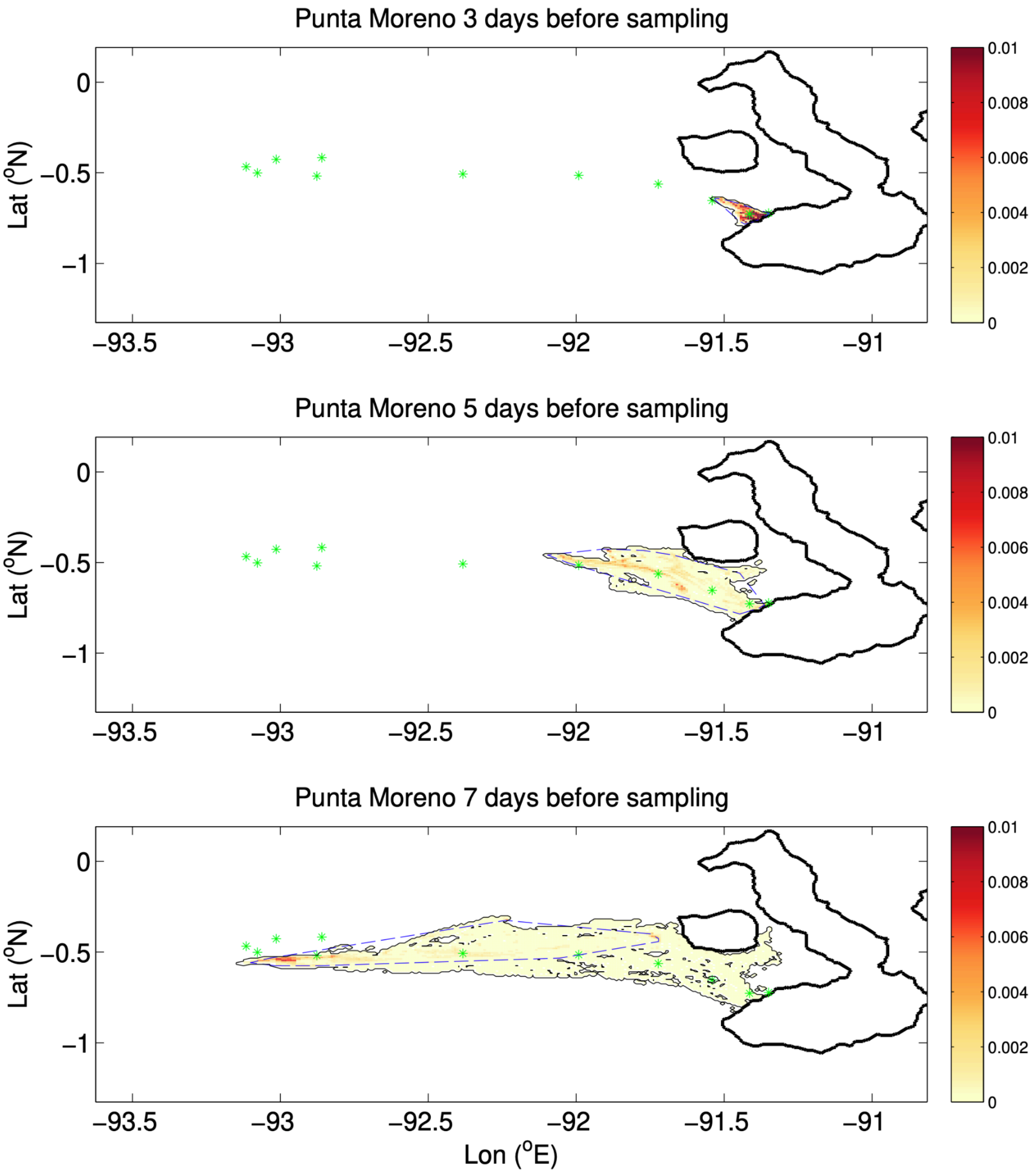

**Fig. S8.** Map of Isabela Island depicting particle backtracking from the ‘Punta Moreno’ site over 3 days (top), 5 days (middle), and 7 days (bottom) prior to the sampling date. Green asterisks indicate the daily particle centroids, with all days displayed across each panel. The red-yellow shading represents the spatial distribution of particle release proportion, as indicated by the color scale on the right. The dashed blue line outlines the area used for calculating the particle surface area metric.

**Table S1.** Sampling site metadata detailing nearest island, co-ordinates, bioregion, sampling date (DD/MM/YYYY), and sampling depth in meters for the shallow and deep samples.

| SiteID | Location | Island | latitude | longitude | Bioregion | SamplingDate | DepthMeters | DepthMeters |
|---|---|---|---|---|---|---|---|---|
| BAR | Barahona Beach | Isabela | -1.03875 | -91.156367 | Western | 30/09/2018 | 3.7 | 1.8 |
| CDOU | Cabo Douglas | Fernandina | -0.302817 | -91.651733 | Western | 03/10/2018 | 3.7 | 0.9 |
| CHAM | Cabo Hammond | Fernandina | -0.469583 | -91.611183 | Western | 02/10/2018 | 5.5 | 0.9 |
| CMAR | Cabo Marshall | Isabela | -0.001333 | -91.22385 | CSouthEastern | 05/10/2018 | 5.4 | 0.9 |
| CORA | Salvaje de Corazón | Genovesa | 0.311317 | -89.97515 | Northern | 08/10/2018 | 5.5 | 0.9 |
| CUEV | Las Cuevas | Floreana | -1.25335 | -90.3741 | CSouthEastern | 12/10/2018 | 8.5 | 0.9 |
| DAPH | Daphne Mayor | Seymour | -0.4258 | -90.372117 | CSouthEastern | 10/10/2018 | 9.8 | 0.9 |
| EGAS | Cerca a Puerto Egas | Santiago | -0.2596 | -90.868783 | CSouthEastern | 09/10/2018 | 21.6 | 0.9 |
| ELI | Marielas-Bahia Elizabeth | Isabela | -0.59505 | -91.0859 | Elizabeth | 01/10/2018 | 4.9 | 0.9 |
| GARD | Bahía Gardner | Española | -1.365467 | -89.639317 | CSouthEastern | 13/10/2018 | 5.8 | 0.9 |
| PCAL | Punta Calle | Marchena | 0.286433 | -90.501817 | Northern | 07/10/2018 | 6.4 | 0.9 |
| PESP | Punta Espinosa | Fernandina | -0.272283 | -91.435383 | Western | 03/10/2018 | 8.5 | 0.9 |
| PIN | Cabo Ibbetson | Pinta | 0.603883 | -90.789833 | Northern | 06/10/2018 | 7.6 | 0.9 |
| PLAZ | Islas Plaza | Santa Cruz | -0.567567 | -90.173617 | CSouthEastern | 11/10/2018 | 5.5 | 0.9 |
| PMAN | Punta Mangle | Fernandina | -0.458567 | -91.390833 | Elizabeth | 02/10/2018 | 3.5 | 0.9 |
| PMOR | Punta Moreno | Isabela | -0.7223 | -91.345067 | Western | 01/10/2018 | 6.7 | 0.9 |
| PROJ | Playa Roja | Rábida | -0.40895 | -90.715533 | CSouthEastern | 09/10/2018 | 18.6 | 0.9 |
| PVIC | Punta Vicente Roca | Isabela | -0.045367 | -91.543017 | Western | 04/10/2018 | 7 | 0.9 |
| RED | Roca Redonda | Isabela | 0.2706 | -91.626483 | Northern | 04/10/2018 | 8.2 | 0.9 |
| SOMB | Sombrero Chino | Santiago | -0.350817 | -90.5685 | CSouthEastern | 10/10/2018 | 7.6 | 0.9 |
| STAFE | Bahía Santa fe | Santa Fe | -0.799483 | -90.04745 | CSouthEastern | 11/10/2018 | 6.1 | 0.9 |
| SUAR | Punta Suárez | Española | -1.350567 | -89.710433 | CSouthEastern | 13/10/2018 | 11 | 0.9 |
| TOR | Isla Tortuga | Isabela | -1.011717 | -90.8767 | Western | 30/09/2018 | 30.5 | 1.8 |

**Table S2.** Outputs from beta diversity statistical models testing for significant within group dispersion (PERMDISP procedure) and between group differences (PERMANOVA) for bioregion differences from eDNA metabarcoding data. The compared bioregions are shown in the leftmost column with the outputs from the PERMDISP procedure in subsequent columns and the results from the PERMANOVA procedure in the last six columns. Significant results (α = 0.05) are shown in bold.

| | Beta Dispersion (PERMDISP) | | | | PERMANOVA | | | | | |
|---|---|---|---|---|---|---|---|---|---|---|
| | diff | lwr | upr | P value | SumofSqs | MeanSquares | F.model | R2 | P value | Adjusted P |
| Elizabeth-CSouthEastern | -0.1393 | -0.2407 | -0.0379 | **0.0053** | 0.4538791 | 0.4539 | 2.2871 | 0.1861 | 0.0134 | **0.0201** |
| Northern-CSouthEastern | -0.0448 | -0.1222 | 0.0327 | 0.3888 | 0.3228144 | 0.3228 | 1.6053 | 0.1180 | 0.0009 | **0.0027** |
| Western-CSouthEastern | 0.0126 | -0.0520 | 0.0771 | 0.9461 | 0.5146647 | 0.5147 | 2.4282 | 0.1393 | 0.0003 | **0.0018** |
| Northern-Elizabeth | 0.0945 | -0.0189 | 0.2079 | 0.1232 | 0.4814028 | 0.4814 | 2.5428 | 0.3886 | 0.0667 | 0.0800 |
| Western-Elizabeth | 0.1518 | 0.0469 | 0.2568 | **0.0034** | 0.2823101 | 0.2823 | 1.2972 | 0.1563 | 0.1093 | 0.1093 |
| Western-Northern | 0.0573 | -0.0247 | 0.1394 | 0.2358 | 0.5056828 | 0.5057 | 2.3314 | 0.2057 | 0.0027 | **0.0054** |

**Table S3.** Generalized least squares linear regression model outputs from models predicting eDNA-derived asymmetric Jaccard dissimilarities from geographic distance (m),oceanographic resistance ($ms^{-1}$), and temperature difference (°C).

| **Model Summary (unscaled variables)** | |
|---|---|
| AIC | -1132.221 |
| BIC | -1106.91 |
| LogLink | 572.1106 |
| rho | 0.1402854 |

| **Variable** | **Value** | **Std.Error** | **t-value** | **p-value** |
|---|---|---|---|---|
| (Intercept) | 0.4537249 | 0.015116927 | 30.0144 | <0.0001 |
| GeographicDistance | 0.0000004 | 0.000000049 | 9.24 | <0.0001 |
| TempDistance | 0.0060691 | 0.001352736 | 4.4866 | <0.0001 |
| OceanResistance | -0.0883384 | 0.023642891 | -3.7364 | 0.0002 |

| **Model Summary (scaled variables)** | |
|---|---|
| AIC | -1152.324 |
| BIC | -1127.012 |
| LogLink | 582.1619 |
| rho | 0.1402854 |

| **Variable** | **Value** | **Std.Error** | **t-value** | **p-value** |
|---|---|---|---|---|
| (Intercept) | 0.5190285 | 0.013360535 | 38.8479 | <0.0001 |
| GeographicDistance_scaled | 0.0318613 | 0.003448194 | 9.24 | <0.0001 |
| TempDistance_scaled | 0.014507 | 0.003233432 | 4.4866 | <0.0001 |
| OceanResistance_scaled | -0.0120814 | 0.003233458 | -3.7364 | 0.0002 |

**Table S4.** Generalized least squares linear regression model outputs from models predicting eDNA-derived asymmetric Jaccard dissimilarities from geographic distance (m),oceanographic resistance ($ms^{-1}$), Lagrangian minimum drift days and temperature difference (°C).

| **Model Summary (unscaled variables)** | |
|---|---|
| AIC | -514.9722 |
| BIC | -490.6663 |
| LogLik | 264.4861 |
| rho | 0.1422763 |

| **Variable** | **Value** | **Std.Error** | **t-value** | **p-value** |
|---|---|---|---|---|
| (Intercept) | 0.4663642 | 0.0158388 | 29.444409 | <0.0001 |
| GeographicDistance | 0.0000002 | 0.00000009 | 2.364411 | 0.0189 |
| TempDistance | 0.0145419 | 0.00272552 | 5.335464 | <0.0001 |
| OceanResistance | -0.063329 | 0.03603917 | -1.757227 | 0.0802 |
| LagranMinimum | 0.0009499 | 0.00049352 | 1.924772 | 0.0555 |

| **Model Summary (scaled variables)** | |
|---|---|
| AIC | -539.4205 |
| BIC | -515.1146 |
| LogLik | 276.7102 |
| rho | 0.1422763 |

| **Variable** | **Value** | **Std.Error** | **t-value** | **p-value** |
|---|---|---|---|---|
| (Intercept) | 0.5031377 | 0.013205536 | 38.10051 | <0.0001 |
| GeographicDistance_scaled | 0.0136613 | 0.005777906 | 2.36441 | 0.0189 |
| TempDistance_scaled | 0.0268296 | 0.005028546 | 5.33546 | <0.0001 |
| OceanResistance_scaled | -0.0092214 | 0.005247672 | -1.75723 | 0.0802 |
| LagranMinimum_scaled | 0.0111216 | 0.005778141 | 1.92477 | 0.0555 |
